# Persistence of ambigrammatic narnaviruses requires translation of the reverse open reading frame

**DOI:** 10.1101/2020.12.18.423567

**Authors:** Hanna Retallack, Katerina D. Popova, Matthew T. Laurie, Sara Sunshine, Joseph L. DeRisi

**Author notes:** Address correspondence to Joseph L. DeRisi.

## Abstract

Narnaviruses are RNA viruses detected in diverse fungi, plants, protists, arthropods and nematodes. Though initially described as simple single-gene non-segmented viruses encoding RNA-dependent RNA polymerase (RdRp), a subset of narnaviruses referred to as “ambigrammatic” harbor a unique genomic configuration consisting of overlapping open reading frames (ORFs) encoded on opposite strands. Phylogenetic analysis supports selection to maintain this unusual genome organization, but functional investigations are lacking. Here, we establish the mosquito-infecting Culex narnavirus 1 (CxNV1) as a model to investigate the functional role of overlapping ORFs in narnavirus replication. In CxNV1, a reverse ORF without homology to known proteins covers nearly the entire 3.2 kb segment encoding the RdRp. Additionally, two opposing and nearly completely overlapping novel ORFs are found on the second putative CxNV1 segment, the 0.8 kb “Robin” RNA. We developed a system to launch CxNV1 in a naïve mosquito cell line, then showed that functional RdRp is required for persistence of both segments, and an intact reverse ORF is required on the RdRp segment for persistence. Mass spectrometry of persistently CxNV1-infected cells provided evidence for translation of this reverse ORF. Finally, ribosome profiling yielded a striking pattern of footprints for all four CxNV1 RNA strands that was distinct from actively-translating ribosomes on host mRNA or co-infecting RNA viruses. Taken together, these data raise the possibility that the process of translation itself is important for persistence of ambigrammatic narnaviruses, potentially by protecting viral RNA with ribosomes, thus suggesting a heretofore undescribed viral tactic for replication and transmission.

**IMPORTANCE:** Fundamental to our understanding of RNA viruses is a description of which strand(s) of RNA are transmitted as the viral genome, relative to which encode the viral proteins. Ambigrammatic narnaviruses break the mold. These viruses, found broadly in fungi, plants, and insects, have the unique feature of two overlapping genes encoded on opposite strands, comprising nearly the full length of the viral genome. Such extensive overlap is not seen in other RNA viruses, and comes at the cost of reduced evolutionary flexibility in the sequence. The present study is motivated by investigating the benefits which balance that cost. We show for the first time a functional requirement for the ambigrammatic genome configuration in Culex narnavirus 1, which suggests a model for how translation of both strands might benefit this virus. Our work highlights a new blueprint for viral persistence, distinct from strategies defined by canonical definitions of the coding strand.

## INTRODUCTION

Narnaviruses are RNA viruses found broadly in eukaryotic hosts including fungi, plants, protists, and arthropods (1). The narnavirus RNA-dependent RNA polymerase (RdRp) is most closely related to mitoviruses and ourmiaviruses, and more distantly to the bacteriophage leviviruses, all of which are positive sense single-stranded RNA viruses (+ssRNA) (2). The canonical narnavirus genome is simple: a single-stranded RNA 2.3-3.6 kb in length, encoding an RdRp (2). Double-stranded RNA (dsRNA) forms can be isolated, although it is unclear whether these represent by-products of RNA extraction, *bona fide* replication intermediates, translation templates, or transmissible units, or some combination of the above (3, 4). Recently, putative second segments have been identified for two narnaviruses with likely protist hosts, and in the arthropod-infecting Culex narnavirus 1 (CxNV1) (5–7). The function of proteins encoded by these segments is unknown, and their association with the RdRp segment is thus far only correlative. In many cases, narnavirus infection does not produce an observable phenotype and persists non-pathogenically, though examples exist where the narnavirus alters its host’s biology (8). From biochemical studies of narnaviruses in *Saccharomyces cerevisiae* and in nematodes, transmission is thought to occur via cytoplasmic inheritance during horizontal transfer in yeast or cell division including animal germ lines, without any extracellular virion (9, 10). As described so far, narnaviruses would appear to exhibit a straight-forward replication cycle. However, additional complexity in the genome organization of some narnaviruses may provide insight into a unique host-virus relationship that has not yet been described for any virus.

A subset of narnavirus genomes have a surprisingly complex feature. In particular, an additional uninterrupted open reading frame (ORF) is found in the reverse direction overlapping the RdRp ORF and spanning almost the entire length of the segment. Such narnaviruses have hence been described as “ambigrammatic,” as both the forward and reverse ORFs have the potential to be translated. This is achieved by avoidance of the codons UUA, CUA, UCA, which encode stops in the –0 frame (1, 11). The extent of overlap in ambigrammatic narnaviruses is unprecedented in the catalog of viral configurations.

Overlapping ORFs are common among viruses. Advantages include compact genomes, novel genes, and translation regulation, at the expense of evolutionary constraints imposed on the sequence by coding in two frames instead of one (12). Notably, amibgrammatic narnaviruses are distinct from other RNA viruses because the ORF overlap is both extensive (>3 kb comprising >95 % of the genome) and anti-parallel, aka pairing forward and reverse frames (1, 13, 14). Intriguingly, although ambigrammatic narnaviruses are diverse with just 27 % mean pairwise amino acid identity in their RdRps, the presence of a reverse ORF (rORF) is conserved in an apparently all-or-none fashion (1), and suggests a potential selection to retain the rORF. To our knowledge, the role of the ambigrammatic feature has not been investigated experimentally and its function is wholly unknown.

These compelling observations motivate our investigation of CxNV1, an ambigrammatic narnavirus found in wild-caught mosquitoes. Its 3.2 kb RdRp-encoding RNA is associated with a second RNA, the 0.8 kb “Robin” segment. Both segments are ambigrammatic. Here, we analyzed the CxNV1 genome in a persistently-infected cell line, developed a launching system to test the requirement for the rORF, examined rORF translation by mass spectrometry, and interrogated the association of ribosomes with each viral RNA by ribosome profiling.

## RESULTS

To refine the genome of a model ambigrammatic narnavirus, we first characterized the complete genome sequence of CxNV1 found in the CT cell line derived from embryos of the *Culex tarsalis* mosquito (Figure 1, Figure S1, Table S1). An end-specific adapter-ligation sequencing approach revealed complementary ends on each CxNV1 segment, beginning 5′-GGGG and ending CCCC-3′ with a 21 nt 3′ hairpin (Figure 1A and B, Figure S2, Tables S2, S3). These features are conserved among narnaviruses (15). Overlapping ORFs on opposite strands extend nearly the full length of each segment. With respect to these shared genomic features, and to further investigate the association between the segments, the antiviral host cell response to each segment was assessed by reanalysis of small RNA sequencing data derived from the same CT cell line, as published by Goertz et al. (16). Abundant small RNAs aligning to both strands of the previously undetected CxNV1 Robin segment were present with a length mode of 21 nt, indicative of a small interfering RNA (siRNA) response similar to that against the RdRp segment (Figure S3). The copy number ratio of Robin to RdRp segments was calculated at ~3.8, based on length-normalized read abundance in meta-transcriptomic next-generation sequencing (mNGS) libraries.

**Figure 1.**
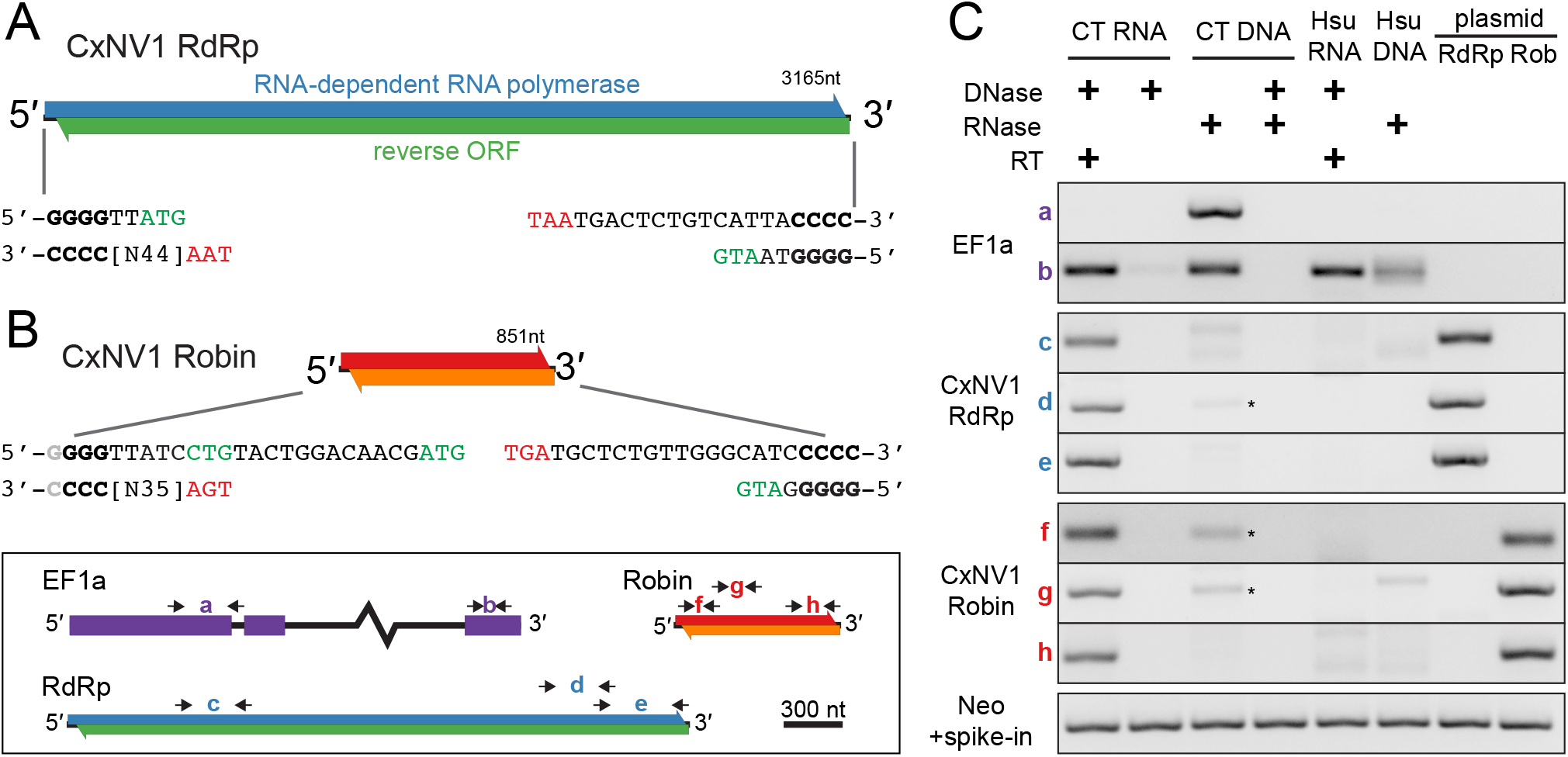
Characterization of CxNV1 in CT cell line. **(A and B)** Schematic of forward and reverse ORFs on the larger segment encoding the RdRp (A), and on the smaller segment, Robin (B). Consensus end sequences shown, with start codons in green and stop codons in red. Light grey nucleotide at 5’ end of Robin indicates similar frequencies observed for 5’-GGGG and 5’-GGG. **(C)** RT-PCR on RNA and DNA from CT and Hsu cells, or on plasmids containing full-length clones of CxNV1 RdRp or Robin, treated with nucleases as indicated, with or without reverse transcriptase (RT) in the reaction. Key for primer pair locations shown in bottom left of figure. Intronic primer specific to *C. tarsalis* in EF1a (primer pair a) indicates DNA recovery (absent from *C. quinquefasciatus* Hsu cells due to intronic sequence variation). Asterisks indicate faint bands at expected size in CT DNA. For reactions in the lowest row, a plasmid containing the neomycin (Neo) gene was spiked in and amplified using Neo-specific primers, to verify that PCR amplification was not impeded by prior DNase treatment of CT/Hsu input.

The unusual layout of ORFs raised the possibility of alternate RNA forms such as circular, sub-genomic, or multiple defective viral RNAs. To address this question, we carefully examined the coverage profile of mNGS data and verified continuity with Sanger sequencing of >1 kb PCR products along the genome. For CxNV1, no evidence for alternate RNA forms was found. In contrast, for Calbertado virus (CALV), another persistent coinfection in CT cells, mNGS clearly showed a large increase in the coverage of the 3′UTR (Figure S4) consistent with known sub-genomic flaviviral RNAs (17). In summary, the RNA forms of both the RdRp and Robin segments of CxNV1 exhibit overlapping ORFs and complementary ends, are full-length linear RNAs, and are targeted by similar antiviral responses.

Next, we sought to determine whether DNA forms exist for CxNV1, which belongs to a family classified as +ssRNA viruses, but infects insect cells which commonly endogenize viral sequences (18). As expected, reverse transcription PCR (RT-PCR) yielded facile amplification of multiple products derived from both CxNV1 RdRp and Robin segments (Figure 1C, Figure S5). In contrast, amplification of CxNV1 from RNase treated DNA was very inefficient, whereas amplification of the host EF1a genomic locus was robust as expected. In all cases, Sanger sequencing of amplification products confirmed on-target product. Variants in the RNA mNGS data were then analyzed. Host genes such as EF1a and GAPDH contained single nucleotide variants (SNVs) at frequencies ranging from 0.45 to 0.50 which were phased as expected for two alleles derived from a diploid genome. In contrast, the SNVs observed in CxNV1 had frequencies far outside the bi-allelic range (<0.05-0.32), included sites with >2 variants, and were not phased (Table S4). Finally, the relative abundance of positive and negative strands was calculated and compared to characteristic ratios for host transcripts or viral transcripts from +ssRNA, dsRNA, or −ssRNA viruses. The relative abundance of positive to negative strand was ~70 for the RdRp segment, and ~145 for the Robin segment, similar to ratios for CxNV1 found in wild-caught mosquitoes by Batson et al. (7), and in the range of other +ssRNA and putative dsRNA viruses (Figure S6).

Overall, these analyses support the bi-segmented nature of CxNV1 as an RNA virus with substantial negative strand contribution. Importantly, the complete and accurate genomes allowed us to attempt to launch the virus in uninfected cells.

### Requirement of reverse ORF

As previous inferences concerning CxNV1 biology have relied on interpreting sequencing data alone, we developed a plasmid-based launch system to introduce CxNV1 into naïve cells in order to test the functional requirement for each segment and their rORFs. Previously, narnaviruses have been launched by introducing the positive-sense RNA of the 20S RNA virus in *Saccharomyces cerevisiae* (19). Here, plasmids expressing full-length positive-sense CxNV1 were transfected into the narnavirus-free Hsu cell line derived from adult ovaries of the *Culex quinquefasciatus* mosquito (Figure 2). After allowing plasmid loss through cell division, cells were counter-selected by FACS to minimize expression from residual DNA. Although some lingering DNA was detectable, RT-PCR showed that the RdRp segment robustly persisted as RNA at least as far as nine weeks post-transfection with regular cell passaging (Figure 2A and B, Figures S7, S8). The Robin segment only persisted when co-transfected with RdRp, and both segments were undetectable when the GDD domain in the RdRp was mutated to render the polymerase inactive (Figure 2A and B). These results demonstrate the operation of a CxNV1 launch platform which enables manipulation of the viral sequences.

**Figure 2.**
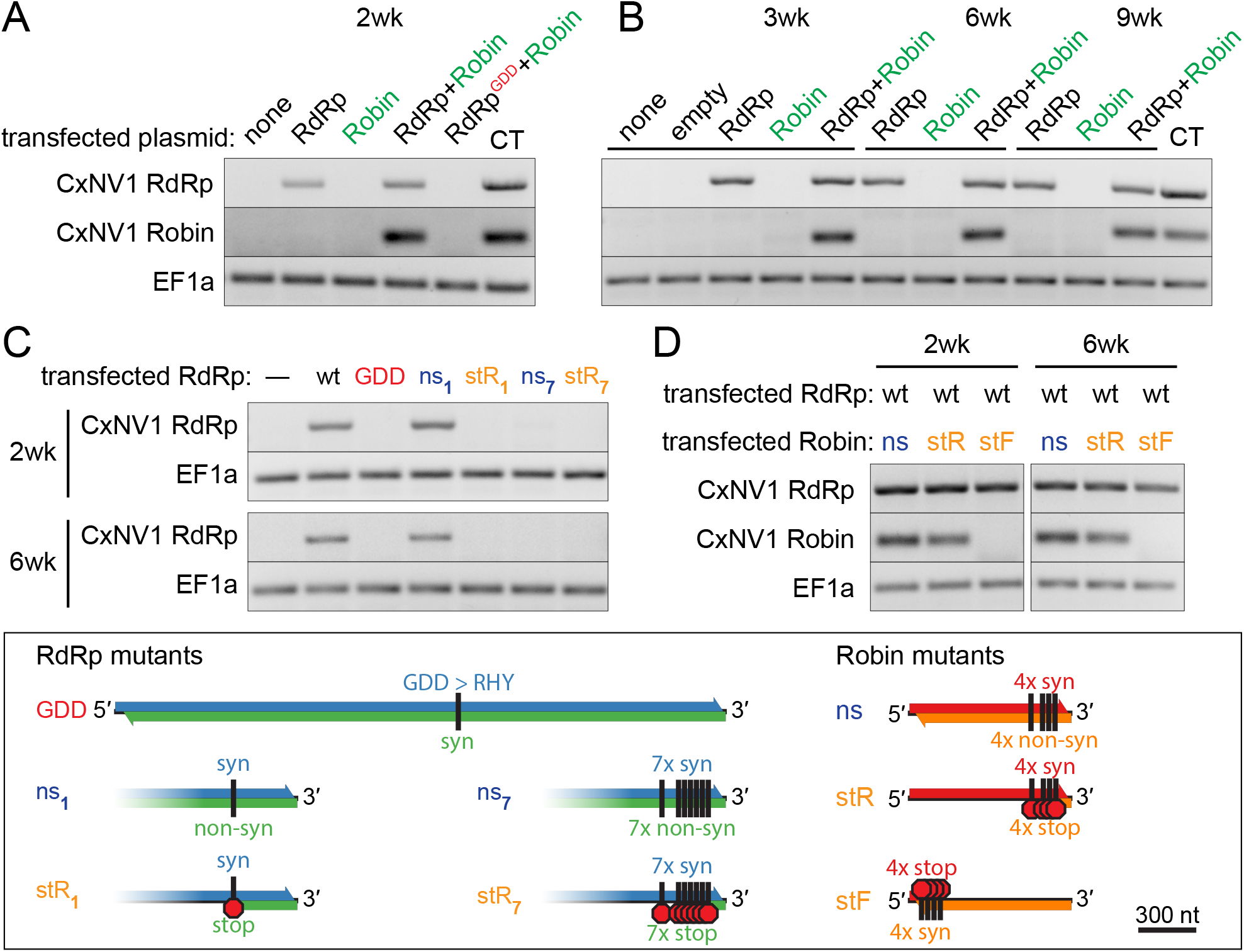
CxNV1 persistence depends on GDD domain and reverse ORF in RdRp. **(A)** RT-PCR targeting CxNV1 RdRp, Robin, or EF1a RNA at 2 weeks post-transfection of Hsu cells with indicated plasmid and sort/counter-sort (see Methods), or CT cells as positive control. Plasmids drive expression of full-length positive-strand viral segments, either wildtype RdRp, inactive mutant RdRp (GDD), and/or wildtype Robin. **(B)** RT-PCR as in (A), for cells collected at 3, 6, and 9 weeks post-transfection. (**C and D)** RT-PCR as in (A), at 2 and 6 weeks post-transfection of Hsu cells with indicated plasmids, wildtype (wt) or mutants diagrammed in key below. **Key:** Mutant RdRp constructs: mutations eliminating conserved motif required for polymerase activity (GDD); mutations introducing stop codons in the reverse ORF while remaining synonymous in the forward ORF (stR); mutations introducing non-synonymous changes in the reverse ORF while remaining synonymous in the forward ORF all at the same nucleotide positions as stR (ns). Subscript (1 or 7) indicates number of codons mutated. Mutant Robin constructs: ns and stR mutants as for RdRp; mutations introducing stop codons beginning at 14 codons from the predicted start site of the forward ORF while remaining synonymous in the reverse ORF (stF). All experiments were performed in biological triplicate, with representative results shown here.

Using this launch system for CxNV1 infection in cell culture, we next dissected the requirement for intact rORFs in the CxNV1 lifecycle. First, mutations were introduced to interfere with the rORF while only introducing synonymous mutations on the opposing strand (Table S5). While a single non-synonymous mutation in the rORF opposing RdRp had no effect (“ns1”), the introduction of a single stop codon in the rORF at the same position (“stR1”) resulted in loss of the RNA by two weeks post-transfection (Figure 2C). Likewise, a more dramatic change introducing seven stop codons in the rORF (“stR7”) also resulted in loss of the RNA. Interestingly, introduction of seven non-synonymous mutations into the rORF opposing RdRp also resulted in loss (“ns7”), suggesting that specific sequence elements in the 3′ region of the RdRp segment are important. Comparable mutations in Robin did not result in loss of the RNA in this assay (Figure 2D). However, the Robin RNA was lost by two weeks when stop mutations were introduced near the beginning of the forward ORF, suggesting either required RNA *cis-*acting features, or that persistence of the Robin RNA requires the protein encoded by its forward ORF. Targeted sequencing of the recovered RNAs showed that all mutations were retained without reversion to wildtype. Together, these data provide evidence that a functional RdRp protein is required for replication of its own RNA segment as well as the Robin RNA segment, and that an uninterrupted rORF is critical for CxNV1 RdRp persistence in this system.

### Interaction of ribosomes with CxNV1 RNA

The requirement for an intact rORF in the essential RdRp segment implies translation of the full-length rORF, or at least a functional interaction of ribosomes with the rORF during CxNV1 infection. To investigate translation in CxNV1 infection, we first asked if rORF protein production could be detected in lysates from CT cells persistently infected with CxNV1. Mass spectrometry demonstrated that peptides from across the entire length of both forward and reverse ORFs on the RdRp segment were indeed present (Figure 3A, Table S6), confirming that translation occurs on both strands of the RdRp segment.

**Figure 3.**
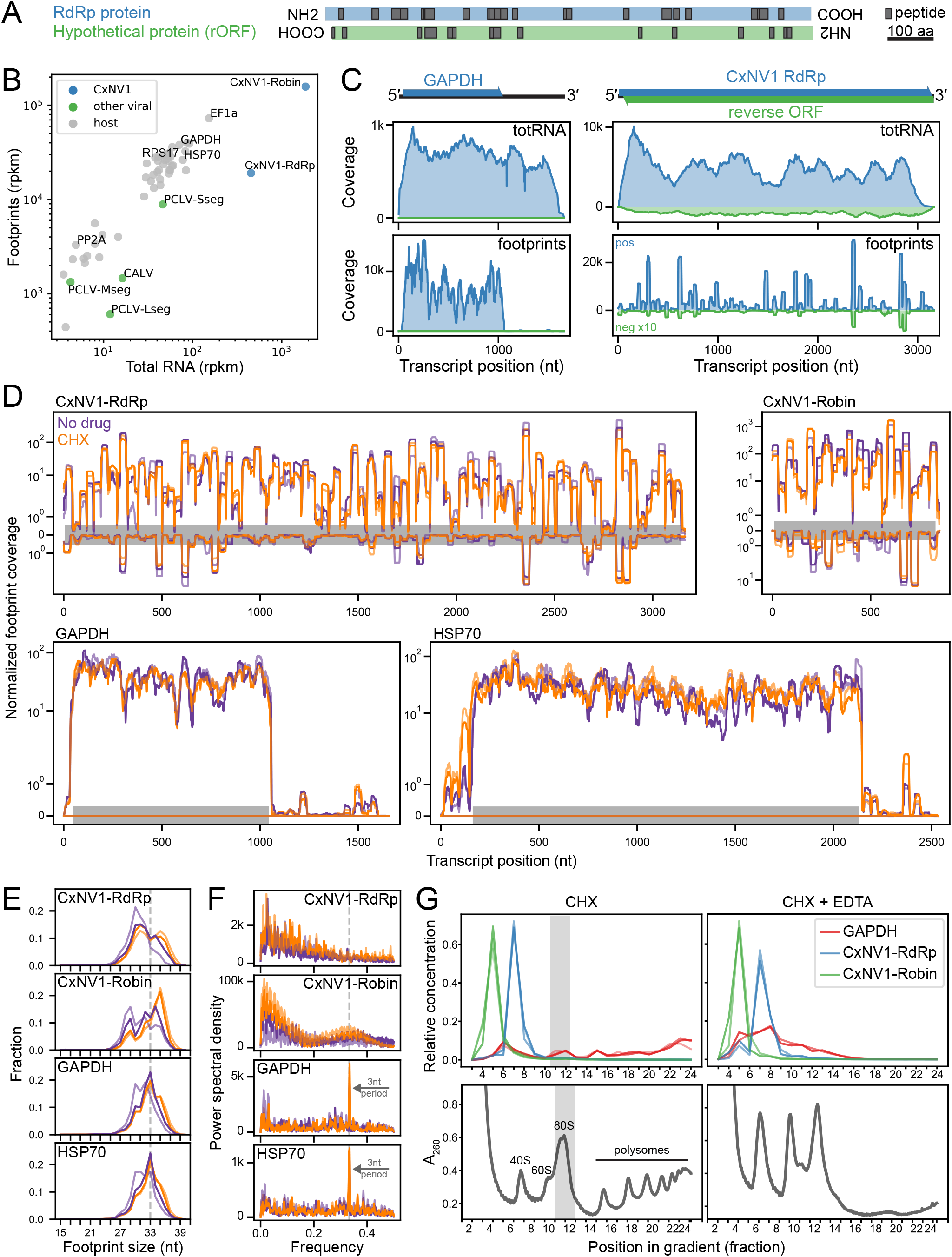
Investigation of ribosome interaction with each CxNV1 RNA strand and segment. **(A)** Alignment of peptides (in black) found by LC-MS/MS of CT cell lysates to the amino acid sequence for the predicted RdRp and Hypothetical proteins encoded by the forward and reverse ORFs, respectively, of the CxNV1 RdRp RNA segment. Displayed according to ORF layout in the genome. **(B)** Relative abundance of selected host transcripts and persistent viruses including CxNV1 in ribosome profiling (footprints) vs. total RNA sequencing libraries. Data shown for positive strand, within-cds densities, CHX-rep1 sample. **(C)** Read coverage of GAPDH and CxNV1-RdRp, for ribosome profiling (footprints) and total RNA sequencing libraries. Reads mapping to negative strand in green, shown with 10-fold y-axis magnification compared to positive strand reads in blue for visualization purposes. Data shown for CHX-rep1 sample. **(D)** Ribosome profiling read coverage of each CxNV1 segment and two host transcripts. Data shown for two replicates of each condition: no drug (purple), CHX (orange), using normalized values and displayed on a log10 scale. Grey bar indicates cds position on positive strand. Reads mapping to negative strand shown inverted. **(E)** Length distribution of footprints for each CxNV1 segment and two host transcripts in ribosome profiling sequencing library. Color indicates condition as in (D). **(F)** Analysis of periodicity in footprints for each CxNV1 segment and two host transcripts. Ribosome densities from 5’ mapping were analyzed using Welch’s method to estimate the power spectral density at each frequency. Frequency of ~0.33 (grey dashed line) corresponds to a period of 3nt. Color indicates condition as in (D). **(G)** Polysome profiling of CT cells treated with CHX (left) or CHX+EDTA (right). Absorbance quantifying total RNA during fractionation of a single gradient for each condition shown in lower panels. Relative concentrations of CxNV1-RdRp, CxNV1-Robin, and GAPDH RNA determined by RT-qPCR in upper panel, with replicates of independent gradients (n=3 for CHX, n=2 for CHX-EDTA). Grey shading indicates monosome peak.

Next, ribosome profiling was performed to explore the translational landscape of CxNV1 RNAs, with a focus on whether ribosomes engaged in typical active translation of CxNV1 ORFs akin to that of host mRNAs, or alternate modes of interaction given the unusual ORF layout in this virus. Ribosome profiling libraries were prepared in biological duplicate from the micrococcal nuclease (MNase)-digested monosome fraction of CT cells with either cycloheximide (CHX) or no treatment (no-drug) before lysis (Table S7). RNAseq libraries were prepared from total RNA (totRNA) of the same samples using a mNGS workflow. Apart from a slight increase in footprint coverage near the 5′ end of some ORFs in the CHX libraries compared to no-drug, the additional pre-treatment had little effect. Thus, the four libraries were considered as near-replicates for the remainder of the analyses.

A selection of 48 host transcripts were *de novo* assembled for the following analyses, as no genome is currently available for *Culex tarsalis* (Table S8, see Methods). Among these host transcripts, a consistent ratio of footprint to total RNA reads was observed (Figure 3B, Figure S9), with footprint reads aligning within the coding sequences (CDS) on the positive-sense RNA as expected (Figure 3C and D, Figures S10).

For CxNV1, footprints were observed on both strands, at a strand ratio similar to that observed in the total RNA libraries (Figure S11). However, footprints on CxNV1 showed several unexpected features. Firstly, the abundance of footprints relative to total RNA abundance was substantially lower for CxNV1 than for host transcripts (Figure 3B, Figure S9). Secondly, the footprints were heavily concentrated at specific positions resulting in an unusual profile (Figure 3C). This “plateau pattern” in coverage was observed on both CxNV1 segments, in all four libraries, in contrast to the more uniform coverage of typical footprint distributions along host genes such as GAPDH and HSP70 (Figure 3D, Figure S12). Thirdly, the lengths of footprints mapping to CxNV1 were broadly distributed from 27-37 nt, and not enriched in the 33 nt size that dominated host genes (Figure 3E, Figure S13). Fourthly, triplet periodicity, which is expected for normally translating mRNAs, was not observed for CxNV1. Although the imprecision of MNase digestion prevented highly accurate read phasing, fourier transform analysis revealed greatest power at a frequency of 0.33 (period of 3 nt) for most host transcripts, and the nucleotide composition of positions within 20 nt of footprint edges was consistent with codon-based ribosome positioning on host transcripts (Figure 3F, Figures S14, S15A). This periodicity is the expected pattern for translating ribosomes, though weak periodicity has also been observed in *E. coli* ribosome profiling as a consequence of MNase sequence specificity rather than ribosome reading frame (20), examined further below. Taken together, these differences in CxNV1 ribosome profiling point to a potentially distinct type of interaction between both strands of this virus and the host translation machinery.

Given the unprecedented genome organization and striking differences in ribosome profiling data for CxNV1, we performed a series of additional analyses to determine whether technical factors in the experimental approach or biological factors distinguishing CxNV1 from other RNAs could explain our observations. With regard to sequencing, insufficient sampling as a source for CxNV1-specific footprint patterns was ruled out, as an ample number of footprint reads were observed for both CxNV1 segments, and low-abundance host transcripts such as PP2A did not show the plateau pattern. PCR jackpotting was also considered as a confounder. An attempt to remove PCR duplicates by collapsing identical footprint+UMI sequences caused greater depletion of CxNV1-mapping footprints than other genes (Figures S16, S17). However, the extreme concentration of CxNV1 footprint peaks could exceed the diversity in a random 5 nt UMI library. Regardless, the CxNV1-specific plateau pattern was retained after duplicate collapse. Thus, sequencing artefact was insufficient to explain the unusual CxNV1 footprints.

Next, steps in the experimental approach upstream of sequencing were examined as possible explanations for the unusual pattern. Sequence biases inherent to multiple steps in sample preparation, including nuclease digestion, adapter ligation, and circularization, are known to affect footprint recovery (21, 22). For instance, MNase is known to cleave more efficiently 5′ of A and T than of C or G (23), a bias also observed for all transcripts in our data (Figure S15B). To ascertain whether this nucleotide bias was sufficient to account for the CxNV1-specific pattern, a model was developed for footprint distribution based on empirical nucleotide frequencies at the footprint edge and footprint length. No significant difference was seen in the correlation between the observed coverage on CxNV1-RdRp and the model-predicted footprint coverage using either the actual CxNV1-RdRp sequence or a scrambled sequence (Figure S18). This result suggests that nucleotide bias at MNase cut sites alone is unlikely to yield the plateau pattern in CxNV1.

Having found no technical explanation for the unusual CxNV1-specific footprint features, we next addressed biological factors that might distinguish CxNV1 from host mRNAs, testing whether the unusual footprints could be attributable to abundant viral RNA or antiviral pathways, RNA secondary structure, or features at the amino acid level. Footprint data for CxNV1 was first compared to other persistent viruses in the cell line, including CALV and the negative-sense, bi-segmented Phasi Charoen-like virus (PCLV). For CALV and PCLV, relatively uniform footprint coverage was observed across the canonical ORFs with expected drop-offs in the 3′ UTRs (Figures S10). To assess whether the CxNV1-specific footprints represent non-specific interactions with abundant RNAs, footprints on the genomic negative strand of PCLV were examined, as an example of an abundant RNA that is likely not translated. The negative-strand PCLV footprints were less abundant than positive-strand footprints (Figure S11), did not show the CxNV1-specific plateau pattern (Figure S10), and had length distributions that were significantly different from the reference EF1a positive strand distribution (PCLV Lseg and Mseg, Figure S13), suggesting that the negative-strand reads for PCLV represented a different process than CxNV1 footprints. Non-coding host transcripts for additional comparison could not be confidently identified given the lack of existing genome for this species, and incomplete annotation of the nearby *Culex quinquefasciatus* genome. Lastly, the distribution of footprints in our ribosome profiling was related to mapping of small RNAs previously sequenced from the same cell line by comparing coverage profiles. No direct correlation or consistent pattern was observed for CxNV1 (Pearson correlation coefficient (*r*) and p value for RdRp: *r*=0.02, p=2.0E-1; Robin: *r*=-0.14, p=2.8E-5). In summary, the plateau pattern of CxNV1 footprints was not solely attributable to abundant viral RNA or antiviral pathways.

We next focused on RNA secondary structure, which is often critical for ribosome recruitment by viruses that lack a 5′ cap, and may serve other functions in regulating translation or influencing interactions with non-ribosomal RNA-binding proteins (RBPs) that in turn might contribute to library preparation in unanticipated ways (24, 25). To approximate the local RNA structure, the mean free energy (MFE) of predicted folding was calculated using sliding windows along each transcript (see Methods). The resulting secondary structure profile did not reveal large internal hairpins in CxNV1, and did not correlate with the footprint profile (Figure S19). With aligned footprint positions, small local peaks of MFE were observed at 5′ and 3′ footprint edges in CxNV1. This modest trend was more apparent in CxNV1 than host genes and was supported in only some regions by mapping footprint densities onto RNA structures predicted from 150-200 nt stretches. Overall, no strong evidence was found to support internal RNA secondary structure as a contributing factor to the CxNV1-specific footprint profiles.

We then considered features at the amino acid level that may affect ribosome positioning. In yeast, codons encoded by less abundant tRNAs tend to be translated more slowly (26). Due to the imprecision of MNase cutting and the absence of an annotated genome for our dataset, A/P/E-sites could not be confidently assigned and codon-based analyses were not performed. It is also possible that features of the nascent polypeptide chain may affect translation elongation and result in footprint pileup, such as the presence of poly-proline stretches or the charge of ~20-30 residues passing through the eukaryotic ribosome exit tunnel (27–31), although the use of cycloheximide may confound early studies in this area. We thus tested for potential biases in the chemical and physical properties of the predicted positive-strand ORFs on CxNV1 compared to other host and viral proteins. No significant bias in charge, hydrophobicity, or individual amino acids enriched in the 20 residues upstream of approximate P-site positions was found (Figure S20).

Finally, polysome profiling was performed to test whether the CxNV1-specific footprint pattern actually represented a snapshot of a single RNA with many protected footprints or a summation of many RNAs each with one protected footprint. RT-qPCR targeting CxNV1 and GAPDH was performed on fractions from the polysome gradient of undigested RNA, ± EDTA treatment of lysates before gradient separation to release RNA-bound complexes including ribosomes (Figure 3G, Figure S21). GAPDH RNA in untreated lysates was found in fractions corresponding to monosomes and polysomes, and was predictably found in lighter fractions after EDTA treatment. Conversely, the vast majority of CxNV1 RNA was found in fractions lighter than monosomes regardless of EDTA treatment, and thus would not be expected to contribute to footprinting data. Furthermore, these data are not consistent with the idea that the CxNV1 profile represents a summation of many sparsely footprinted RNAs. Instead, they raise the possibility that the observed footprints are derived from a small amount of heavily-protected RNA.

Taken together, these data are consistent with the possibility that the majority of CxNV1 RNAs are not actively translated, yet some small fraction of both strands are bound by ribosomes in an unusual configuration that does not appear to reflect active processivity expected from translating complexes. Technical and biological factors that could influence the footprints such as sequence bias, RNA secondary structure, or amino acid composition, did not account for these observations.

## DISCUSSION

Overlapping genes are important features of many viruses with functions ranging from novel gene creation to regulation of expression. Among RNA viruses without DNA intermediates, the vast majority of ORF overlaps are <1.5 kb and occur in the same direction (12, 13, 32), and translation of overlapping antisense ORFs has never been experimentally demonstrated (1, 14). In this context, the antisense 3.1 kb overlap in CxNV1 is especially striking. Until now, investigation of this ambigrammatic narnavirus has been limited to *in silico* analyses for lack of an experimental system, leaving unanswered questions about potential functions for antisense overlaps and the specific role of the rORF in the narnavirus lifecycle. Here, we leveraged a naturally persistent infection by CxNV1 of CT cells to address unique aspects of CxNV1 biology including the relationship between the two segments, the requirement for the rORF, and the translational landscape of this unconventional virus.

Our data suggest that CxNV1 persists predominantly as RNA in CT cells. While the amount of dsRNA was not explicitly quantified in this study, the over-abundance of positive strand RNA implies that the possible dsRNA fraction may be no more than 1-2% of the total, if it exists. The faint signal of an RNA virus detected in the DNA fraction of infected CT cells is not unprecedented (33). While viral sequences are frequently endogenized in insect cells (18, 34, 35), a bi-allelic pattern of SNV frequency expected for such endogenized sequences was not observed for CxNV1. Taken together, the data are more consistent with non-endogenized viral DNA likely attributable to incidental reverse transcription of the CxNV1 RNA.

The presence of a second segment associated with CxNV1 (the “Robin” segment) is a relatively new and unexplored finding. Our data suggest that CxNV1 RdRp is required for replication of the Robin RNA. While the RdRp segment persisted without Robin in cell culture, this RdRp-only state has not been observed for CxNV1 in wild-caught mosquitoes (7). It is possible that Robin provides a fitness benefit that is only apparent in the context of the complete organism, or on a time scale not reproduced by cell culture. Robin may also have a niche role for CxNV1 specifically, as it is unclear whether comparable segments exist in other ambigrammatic narnaviruses, and the CxNV1 Robin genome organization is unlike those of non-RdRp segments identified in the distantly related narnaviruses LepseyNLV1, MaRNAV-1 and MaRNAV-2, (5, 6). Additional sequences in the phylogenetic tree are needed to clarify the evolutionary history of the Robin segment and whether its relationship with the RdRp is an example of viral parasitism or symbiosis.

The relationship between the RdRp and Robin segments of CxNV1 may hinge on the shared feature of overlapping ORFs on opposite strands. Our experiments demonstrated that premature truncation of the rORF on the RdRp segment resulted in loss of the RNA, indicating a requirement for the presence of the rORF. As to the importance of the rORF protein sequence, we note that the forward ORFs are considerably more conserved than the rORFs, as shown in a previously published dataset of related CxNV1 sequences (7), where the number of variable positions per 100 amino acids decreases as follows: 47 in the Robin rORF > 40 in the RdRp rORF > 36 in the Robin forward ORF > 13 in RdRp forward ORF encoding the RdRp protein. This order may reflect the importance of the RdRp protein relative to other viral proteins. Alternatively, the lack of rORF sequence conservation may indicate that the act of translation is important rather than a specific protein product.

What, then, is the role of the essential overlapping opposite-sense ORFs? While the rORF appears to be translated, the bulk of viral RNAs appear to be devoid of ribosomes. Assuming that the footprints observed in the ribosome profiling libraries truly indicate ribosome-protected RNA, then the interaction of ribosomes with CxNV1 is unlike typical host RNA translation. We propose a model where a small fraction of CxNV1 RNA is occupied by regions of densely packed ribosomes (Figure 4). This configuration, on each strand of both segments, might result from one or more stall points with ribosome queueing, yielding discrete positions with 30-40 nt spacing. It is unclear whether the CxNV1 proteins are primarily generated from this configuration or from other viral RNAs with more efficient translation, possibly segregated by subcellular compartmentalization. It has long been known that stacking of up to 10 ribosomes can occur (36), and recent attention on disomes suggests that ribosome collisions may be a widespread phenomenon on eukaryotic cellular transcripts (37–39). Our model prompts the question of how CxNV1 might interact with mechanisms for handling ribosome collisions through rescue or degradation (40). The possible causes of ribosome pausing on CxNV1 are also unclear – factors may include RNA secondary structure that cannot be easily predicted from the sequence, in addition to the conserved hairpin at the 3′ end of the positive strand. It also remains unclear how CxNV1 regulates competition between the processes of replication and translation.

**Figure 4.**
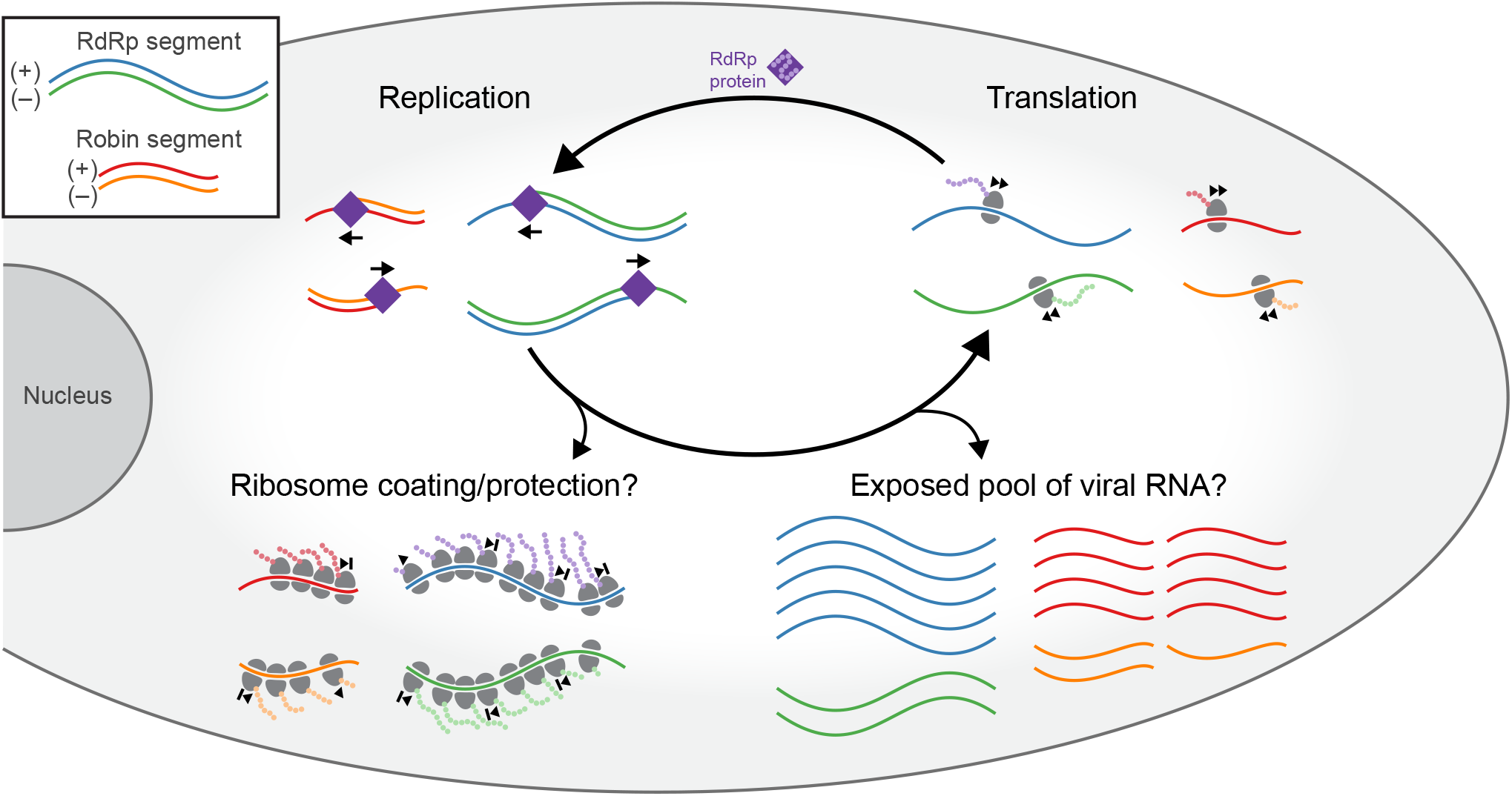
Model of CxNV1 infection. The RdRp segment encodes the viral RNA-dependent RNA polymerase (RdRp), which replicates both the RdRp and Robin segments to generate (+) and (–) strand viral RNA. A small fraction of the total viral RNA may be densely covered in ribosomes. These ribosomes may queue in predictable locations behind paused ribosomes, thus protecting the viral RNA. The remaining pool of viral RNA may be exposed to nucleases. Actively translating ribosomes also produce proteins from the ORFs on each strand of each segment. The role of proteins encoded by the three non-RdRp ORFs is unknown. Ultimately, unconventional interaction of ribosomes with each strand may be more important for the persistence of ambigrammatic narnaviruses than the protein products of translation.

We can only speculate on how ribosome-coating of the viral RNAs might contribute to viral fitness. Perhaps ribosomes fill roles typically performed by nucleocapsid proteins in other viruses, such as protecting the viral RNA from host cell degradation, or maintaining RNA in a polymerase-accessible state. Alternatively, ribosome-coating of the viral RNA could impact cellular ribosome homeostasis in a manner conducive to viral persistence. Finally, the observed ribosome association could have implications for transmission of the CxNV1 RNA, during cell division or elsewise.

Alternative interpretations of the data including technical caveats were considered, but these were found to be insufficient to explain the observations. We considered and rejected artefacts from PCR and library preparation steps, and bias in nucleotide and amino acid sequence. Conceivably, the footprints might represent contaminating RNA or protection by a non-ribosomal RNA-bound protein (RBP) found at the same sucrose density as the monosome peak. Certainly the unusual footprint size distribution for CxNV1-mapping footprints warrants consideration. On one hand, non-conforming size distributions are often used to exclude regions lacking *bona fide* translation activity, as in many non-coding RNAs and 3′ UTRs (41). For unusual-sized footprints mapping to non-coding strands of RNA viruses including influenza and coronaviruses, some groups have hypothesized that the footprints reflect protection by viral nucleoprotein complexes (of note, these protocols isolate footprints using size-exclusion chromatography spin columns or sucrose cushion rather than purifying monosomes via sucrose gradient) (42–45). On the other hand, footprint size can vary according to ribosome conformation, which may in turn be determined by the stage of translation and/or stalled or collided state that is captured at the time of digestion (46). Our technical approach may have captured RNA protected by single ribosomes in unusual conformations. Disomes were likely excluded by the steps of monosome purification and RNA fragment size selection of ~26-34 nt. For CxNV1, non-conforming footprint sizes could be expected from ribosomes that are stacked or otherwise not translating efficiently. Although non-ribosomal RBP footprints cannot be excluded, our model is more parsimonious in joining the observations of dual-ORF conservation and the footprinting data.

In conclusion, this investigation suggests a previously unappreciated strategy among viruses, whereby a small fraction of the RNA from this unique narnavirus exists in a state characterized by bound, densely packed, non-processive ribosomes. This may represent a novel mechanism by which an RNA virus can co-opt the host translational machinery to facilitate persistent infection and transmission in the absence of an intact virion and extracellular phase.

## MATERIALS AND METHODS

### Cell Culture

Established cell lines derived from *Culex tarsalis* embryos (CT) and *Culex quinquefasciatus* ovaries (Hsu) were a generous gift of Aaron Brault (Centers for Disease Control and Prevention, Fort Collins, CO, USA) (47, 48). Cells were grown in Schneider’s Drosophila Medium (Gibco) supplemented with 10 % v/v fetal bovine serum at 28 °C in room air, passaged by scraping, and tested negative for mycoplasma monthly.

### Virus NGS and PCR

To sequence the parental CT cell line, RNA was first extracted using Direct-Zol RNA MicroPrep (Zymo Research). After adding ERCC RNA Spike-In Mix (Life Technologies), the sequencing library was prepared using NEBNext Ultra II Directional RNA Library Prep Kit (NEB), and sequenced with a paired-end 150 bp run on an Illumina NextSeq. The raw reads were submitted to the IDSeq pipeline (v4.6, nt/nr database 2020-02-10) for initial processing including quality filtering, deduplication, host subtraction, de novo assembly, and identification of viral sequences (49). These filtered reads and assembled contigs were used for analyses of the persistent viruses, including positive to negative strand ratios, consensus viral sequences and variant analyses. Bowtie2 (v2.2.4) was used for alignments to correct any assembly errors (50). Independently, raw reads were quality filtered via PRICE Sequence Filter (v1.2, PriceSeqFilter), and used for host analyses including building the host transcriptome reference as described in the Ribosome Profiling section below (51).

Two strategies were used to identify the ends of the CxNV1 RNA: ligation to RNA (for 3′ end), or reverse transcription then ligation to cDNA (for 5′ end). See Table S9 for oligo sequences. For both strategies, the oligo ligated to the unknown end contained a handle TATGCA followed by 0-5 Ns before the Illumina TruSeq Adapter 5 (TSA5) sequence to allow for de-phasing during read1 sequencing (oHR546/547/548/549/550/551). For the RNA ligation method, these ligation oligos were ordered with 5′ phosphorylation and 3′ blocking with C3 Spacer (IDT), pooled in equimolar ratios, pre-adenylated at the 5′ end using Mth RNA Ligase (NEB), then ligated to total RNA extracted from CT cells using 5′ App DNA/RNA ligase (NEB). Next, a TSA5-complementary oligo (oHR552) was used to initiate reverse transcription with SSIII (Thermo Fisher) followed by basic RNA hydrolysis and column purification (DNA Clean and Concentrator, Zymo Research). Then, the regions of interest were amplified and barcoded using nested PCR with Phusion High-Fidelity DNA Polymerase (NEB), an internal primer containing TSA7 and a footprint sequence ~100-150 nt from the predicted end (oHR542/543/544/545, 0.1×), and external primers containing the P5/P7 and i5/i7 Illumina adapter and index sequences.

For the cDNA ligation method, targeted reverse transcription was performed on the extracted CT cell RNA with the footprint oligos oHR542/543/544/545 and SSIII (Thermo Fisher), which is known to add a few non-templated bases after reaching the 5′ end of the RNA with a preference for cytosine (52). The pooled 3′-blocked, 5′-preadenylated oligos oHR546-551 were then ligated to the cDNA using 5′ App DNA/RNA ligase (NEB). Amplification and barcoding PCR was then performed with oligos that annealed to the TSA5 and TSA7 sequences and added i5/i7 and P5/P7 sequences. All reactions were performed according to manufacturer instructions. The final amplicon libraries were size-selected using AMPure XP beads (Beckman Coulter) at 0.9×, then quantified using Qubit dsDNA HS (Thermo Fisher) and size-verified with BioAnalyzer (Agilent). Libraries that failed to enrich for the expected peak size (likely due to non-specific primer binding, and inefficiency of ligating to total RNA before reverse transcription) were dropped at this stage. Final libraries were pooled and sequenced with a paired-end 150 bp run on an Illumina NextSeq.

For analysis of ends libraries, reads were first quality filtered using PriceSeqFilter with flags -rqf 85 0.98 -rnf 90, then trimmed to the read beyond the handle TGCATA. Reads were then aligned to the consensus sequence for the region of CxNV1 well-supported by >4 reads from mNGS sequencing (see above), using Bowtie2 (v2.2.4). Variants in the terminal 8 bases were analyzed to determine the most likely consensus sequence, taking into account potential biases from reverse transcription and ligation, and potential biological variation. RNA folding of terminal sequences was performed using the RNAstructure web server with temperature according to narnavirus host species: 28 °C for insect, 30 °C for yeast (53).

Experiments to determine whether CxNV1 has DNA forms were initiated by extracting RNA from CT cells using Direct-Zol RNA MicroPrep (Zymo Research), and separately, DNA from matched aliquots of CT cells using Quick-DNA Miniprep with Proteinase K (Zymo Research). As a control, RNA and DNA were also extracted from Hsu cells. Extracted RNA was treated with DNase during column purification, and extracted DNA was treated with RNaseA (Thermo Fisher), and/or with DNAse (Invitrogen) as a control. Amplification reactions were performed using KAPA SYBR FAST One-Step qRT-PCR (Roche), with or without reverse transcriptase, on the purified RNA or DNA spiked with a Neo-containing plasmid, with 35 cycles, 60 °C annealing temperature, 30 sec extension, and the following primer pairs: EF1a oHR561/565 (intronic) and oHR672/673 (exonic), CxNV1 RdRp oHR504/509 and oHR502/503 and oHR513/514, CxNV1 Robin oHR653/654 and oHR538/539 and oHR540/541, Neo oHR691/692 (Table S9). PCRs and agarose gel electrophoresis to analyze their products were run in parallel for all conditions. To determine the identity of faint bands at similar size to target products, reactions were re-run with 40 cycles, then bands cloned using Topo TA cloning (Thermo Fisher) and Sanger sequenced.

### Re-analysis of published sequencing data

Small RNA sequencing of CT cells ± infection with West Nile Virus (WNV) was previously performed by Goertz et al. (16). The datasets from NCBI’s sequence read archive (SRA) (accessions SRR8668667 and SRR8668668) were downloaded, adapter sequences trimmed as needed using Trimmomatic (v0.39) with Illumina TruSeq smallRNA adapters and flags ILLUMINACLIP:$adapter_fasta:1:0:0:1 MINLEN:18, quality filtered using PriceSeqFilter with flags -rqf 85 0.98 -rnf 90, then aligned to the combined CT host + virus transcript reference (see Ribosome Profiling Analysis section) using bowtie (v1.2.3) with flags -v 1 -k 1 -m 1.

mNGS of wild-caught mosquitoes in California was previously performed by Batson et al. (7). Quality-filtered reads from the dataset available in the SRA (PRJNA605178) were aligned to the assembled contigs for each sample, for contigs assigned to viral species by the original study authors. The ratio of concordantly aligned read pairs derived from positive (coding) vs. negative strand was calculated. Viral segments with <100 concordantly mapped read-pairs or <3 samples (individual mosquitoes) were discarded from the analysis.

### Plasmids

To clone CxNV1 from CT cell RNA, an iterative process of RT-PCR (Superscript III platinum 1-step RT-PCR, Thermo Fisher) and topo cloning (Invitrogen) of long genomic fragments was performed, ultimately completed with primers oHR594/593, oHR592/586, oHR585/589, and oHR619/620 for 4 fragments of RdRp, and oHR621/622 and oHR623/624 for 2 fragments of Robin, each flanked by BbsI or BsaI cut sites (Table S9). The fragments were assembled into a pUC19 vector using In-Fusion cloning (Takara Bio) with inverse PCR to add ends determined by ligation sequencing method above, ultimately generating plasmids pHR96 (RdRp) and pHR97 (Robin) which contain SmaI-flanked full-length genomic RNAs.

Mutant versions of RdRp and Robin were generated with a combination of PCR, geneblocks (IDT), restriction enzyme and In-Fusion cloning. To control for effects of mutations at the RNA level, mutants with stop codons were compared to mutants with non-synonymous changes at identical nucleotide positions. Unique restriction enzyme sites were introduced to distinguish plasmids easily. Details can be found in Table S5.

To generate plasmids for expression of CxNV1 RNAs in insect cells, a backbone was first created by inserting the T7-HDR (Hepatitis D ribozyme) region from p2RZ (gift from Kristeene Knopp) into pIEX-4 (gift from Wesley Wu), then inserting this cassette into pAc5-STABLE1-Neo (gift from Rosa Barrio # James Sutherland, Addgene plasmid #32425) generating “empty” plasmid pHR106. The RdRp or Robin segments were inserted into this backbone, generating pHR107 and pHR108 respectively, each containing separate Ac5-GFP and hr5/IE1-narnavirus expression constructs. In these plasmids, narnaviral RNA is transcribed with ~72nt upstream of the viral 5′ terminus, and with the Hepatitis D ribozyme to cleave at the viral 3′ terminus. In addition, the GFP was swapped for mCherry in expression plasmids containing CxNV1-Robin.

### Virus Launch

To launch CxNV1 in a naïve cell line, Hsu cells were transfected using X-tremeGENE HP DNA Transfection Reagent (Roche) with plasmid(s) expressing a fluorophore under control of the Ac5 promoter, and the CxNV1 RdRp, Robin, mutants, or empty under control of the IE1 promoter with hr5 enhancer, described above. In initial experiments both segments were in GFP-expressing vectors; subsequent experiments were performed with Robin in a backbone expressing mCherry instead of GFP, showing that when multiple plasmids were transfected concurrently, >95 % of cells received either both or neither plasmid, and yielding no difference in results. As antibiotic-based selection was inefficient, cells were first sorted for successful transfection at 2-6 days post-transfection (fluorophore-positive), then passaged and counter-sorted for loss of plasmid at 10-12 days post-transfection (fluorophore-negative), using a Sony SH800S sorter. At timepoints from 2-9 weeks post-transfection, cells were collected for nucleic acid extraction (including DNase treatment for RNA extractions) and PCR amplification. To determine persistence of viral RNA after loss of plasmid DNA, RT-PCRs were performed with the following primer pairs: oHR672/673 (EF1a), oHR513/514 (CxNV1 RdRp), oHR653/654 (CxNV1 Robin), oHR691/692 (Neo) (Table S9).

### Mass Spectrometry

Lysates were prepared from two 15 cm tissue culture plates of CT cells by lysing in ice-cold RIPA buffer (10 mM Tris-HCl pH 7.4, 1 % v/v Triton X-100, 0.1 % m/v SDS, 140 mM NaCl) with cOmplete EDTA-free protease inhibitor cocktail (Roche), then rotating overhead for 10 min at 4 °C, then centrifuging for 10 min at 16,000 ×g at 4 °C and flash-freezing the supernatant. Protein concentration was measured using the Bradford assay (Bio-Rad). Lysates were diluted in 2× sample buffer (4 % m/v SDS, 20 % v/v glycerol, 120 mM Tris-HCl, 0.02 % m/v bromophenol blue) supplemented with 10 % v/v beta-mercaptoethanol, boiled at 95 °C for 3 min, sheared with 26 G needle, boiled 2 min, centrifuged, then 15 μg loaded onto a NuPAGE 4-12 % Bis-Tris polyacrylamide gel (Invitrogen) and separated by electrophoresis. Gel bands were cut from two regions: 100-150 kD and 25-37 kD and processed via trypsin digestion and LC/MS detailed below. Coomassie stain of parallel gels confirmed adequate separation of the complex lysate. Biological duplicates were performed 10 months apart.

Mass spectrometry was performed by the Vincent J. Coates Proteomics/Mass Spectrometry Laboratory at UC Berkeley. A nano LC column was packed in a 100 μm inner diameter glass capillary with an emitter tip. The column consisted of 10 cm of Polaris c18 5 μm packing material. The column was loaded by use of a pressure bomb and washed extensively with buffer A (see below). The column was then directly coupled to an electrospray ionization source mounted on a Thermo-Fisher LTQ XL linear ion trap mass spectrometer. An Agilent 1200 HPLC equipped with a split line so as to deliver a flow rate of 300 nl/min was used for chromatography. Peptides were eluted with a 90 minus gradient to 60 % B. Buffer A was 5 % acetonitrile/ 0.02 % heptaflurobutyric acid (HBFA); buffer B was 80 % acetonitrile/ 0.02 % HBFA.

Protein identification was done with Integrated Proteomics Pipeline (IP2, Integrated Proteomics Applications, Inc. San Diego, CA) using ProLuCID/Sequest, DTASelect2 and Census (54–57). Tandem mass spectra were extracted into ms1 and ms2 files from raw files using RawExtractor (58). Data was searched against a database consisting of the *Culex quinquefasciatus* and custom viral databases supplemented with sequences of common contaminants. The database was concatenated to a decoy database in which the sequence for each entry in the original database was reversed (59). LTQ data was searched with 3000.0 milli-amu precursor tolerance and the fragment ions were restricted to a 600.0 ppm tolerance. All searches were parallelized and searched on the VJC proteomics cluster. Search space included all fully tryptic peptide candidates with no missed cleavage restrictions. Carbamidomethylation (+57.02146) of cysteine was considered a static modification. Phosphorylation was searched as a variable modification. We required 1 peptide per protein and both trypitic termini for each peptide identification. The ProLuCID search results were assembled and filtered using the DTASelect program (54, 56) with a peptide false discovery rate (FDR) of 0.001 for single peptides and a peptide FDR of 0.005 for additional peptides for the same protein.

### Ribosome Profiling

CT cells were pretreated with 100 μg/mL cycloheximide (CHX, Sigma) at 37 °C for 2 min. Ribosome-protected footprints from cycloheximide-pretreated and no-drug samples were prepared for sequencing as described in a recently updated protocol for ribosome profiling of mammalian cells, with slight modifications (60). Briefly, cells were rapidly harvested at 4 °C in lysis buffer (20 mM Tris-HCl pH 7.4, 150 mM NaCl, 5 mM MgCl_2_, 1 mM DTT, 100 μg/mL CHX, supplemented with 1 % v/v Triton X-100 and 25 U/mL Turbo DNAse I (Thermo Fisher)). Clarified cell lysates were treated with micrococcal nuclease (MNase) to digest RNA not protected by ribosomes. MNase has previously been effective for footprinting insect cells (61), and it has been suggested that RNase I may degrade insect ribosomes (62). In our hands, MNase digestion produced a similar increase in the monosome (80S) fraction in CT cells as RNase T1 (Thermo). 80S ribosomes were isolated by centrifuging lysates through a 10-50 % m/v sucrose gradient at 35,000 rpm for 3 hr at 4 °C with a SW41 rotor on a Beckman L8-60M ultracentrifuge, then collecting the monosome fraction on a BioComp Gradient Station. RNA was then purified from the monosome fractions using a Direct-Zol RNA kit (Zymo Research), then resolved by electrophoresis through a denaturing gel, and the fragments corresponding to ~26-34 bp were extracted.

The 3′ ends of the ribosome footprint RNA fragments were then treated with T4 polynucleotide kinase (NEB) to allow ligation of a pre-adenylated DNA linker with T4 Rnl2(tr) K227Q (NEB). The DNA linker incorporates sample barcodes to enable library multiplexing, as well as unique molecular identifiers (UMIs) to enable removal of duplicated sequences. To separate ligated RNA fragments from unligated DNA linkers, 5′-deadenylase (Epicentre) was used to deadenylate the pre-adenylated linkers, which were then degraded by the 5′-3′ ssDNA exonuclease RecJ (NEB). After rRNA depletion using the Ribo-Zero Gold rRNA removal kit (Illumina), the RNA-DNA hybrid was used as a template for reverse transcription with Superscript III (Thermo), followed by circularization with CircLigase (Epicentre). Finally, PCR of the cDNA circles attached suitable adapters and indices for Illumina Sequencing, quantified and pooled. The library was sequenced with a single-end 50 bp run on an Illumina HiSeq4000.

The corresponding RNA-seq samples were prepared from total RNA of the same cell lysates. RNA was extracted using Direct-Zol RNA MicroPrep (Zymo Research), then libraries prepared using NEBNext Ultra II Directional RNA Library Prep Kit (NEB), and finally sequenced with a paired-end 150 bp run on an Illumina NextSeq.

### Ribosome Profiling Analysis

No genome reference was available for *Culex tarsalis*. The closest species, *Culex quinquefasciatus*, is homologous enough in exonic regions for credible alignments of 150 nt paired-end reads, but we found non-coding regions including introns to be dissimilar enough to make genome-wide alignments unreliable, and annotations somewhat lacking. Instead, we chose to assemble a set of transcripts likely to be well-conserved and spanning from low to high abundance, namely genes for elongation factors, initiation factors, and ribosomal proteins, and a handful of likely single-copy canonical genes, EF1a, GAPDH, HSP70, RPS17, and PP2A. We abandoned genes where the annotations were unclear and which may have multiple copies in the genome. We first extracted mRNA sequences for genes of the classes listed above from the CulPip1.0 genome assembly (NCBI). We then aligned the quality-filtered reads from the 150 nt paired-end sequencing of the parental CT cell line. For transcripts with promising homology, we extended the contig into 5′ and 3′ UTRs using PriceTI (v1.2), then validated by re-mapping paired-end reads, and finally selecting transcripts with unambiguous mapping at >4 read coverage. See Table S8 for Blast results of final host reference transcripts.

The genomes for persistent RNA viruses in the CT cell line were generated by aligning 150 nt paired-end reads to the contigs assembled by Spades in the IDSeq pipeline for error correction and contig extension. Culex narnavirus 1 (CxNV1), Calbertado virus (CALV), and Phasi Charoen-like virus (PCLV) were assembled this way. Anomalies in the alignments and assembly of Flock house virus (FHV) were suspicious for defective viruses or other multiplicity, and so FHV was removed from consideration.

For processing of ribosome profiling data, linker sequences were removed and samples were de-multiplexed using FASTX-clipper and FASTX-barcode-splitter (FASTX-Toolkit v0.0.13). To generate quality and duplication metrics for the entire library, reads were processed with FASTQ-quality-filter with flags -Q33 -q 37 -p 80, then trimmed to remove the 3′ sample barcode (4 nt) and a single base of low quality at the 5′ end (1 nt), then processed with cd-hit-est (v4.8.1) with flag -c 1 to require 100 % identity. For all other analyses, a preliminary alignment of sample barcode- and UMI-trimmed reads to the reference was performed with bowtie (v1.2.3) and flags -v 1 -k 1 -m 1 to allow a single mismatch, then original sequences retrieved, sample barcodes trimmed and quality filtering performed as above. At this stage, reads were optionally collapsed on 100 % identity of the read+UMI using cd-hit-est as above, then UMIs trimmed. From this step forward, parallel pipelines were used for the collapsed and uncollapsed reads. Reads were finally mapped to the reference using bowtie as above, with samtools (v0.1.19) to generate sorted, indexed BAM files.

For the corresponding RNA-seq datasets, reads were first quality-filtered using PriceSeqFilter (v1.2) with flags -rqf 85 0.98 -rnf 90, then mapped to the reference using bowtie2 (v2.2.4) with default parameters. From these BAM files, quantification of reads within the cds on each strand was performed using Bedtools coverage, and coverage at each nt position was determined using Bedtools genomecov with flag -d. Separate mapping to the reference transcripts plus custom *Culex tarsalis* mitochondrial genome (derived from data from Batson et al. (7)) plus ERCCs validated the strand-specificity of library preparation and inclusion of RNA with minimal to no DNA contamination.

Custom scripts were used for further analysis and plotting in Python 3.8, including a combination of plastid (v0.5.1) (63), biopython (v1.77), numpy (v1.19.1), pandas (v1.1.1), scipy (v1.5.2), matplotlib (v3.3.1), and seaborn (v0.10.1). When not otherwise specified, footprint coverage was determined by processing the BAM file via plastid get_count_vectors.py, specifying 27-39 nt footprint length and flags --center --nibble 0 --normalize (distributes the count of 1 read across all positions of the read and normalizes by millions of mapped reads in the sample).

For periodicity analysis, footprints were first mapped to the 5′ end using plastid with -- fiveprime --normalize, then scaled within each refseq+sample pairing to be comparable across samples. Selecting the cds region with 25 nt padding from either end, the power spectral density was estimated using Welch’s method performed by scipy.signal.welch with parameters window= ‘hann’, nperseg=500, noverlap=250, scaling=‘density’, average=‘median’.

For modeling the impact of nucleotide bias on footprint position, we first developed a footprint-generator as follows: choice of 5′ cut site on a given sequence using numpy’s random.default_rng function with probabilities based on the empirical observations of bases found at the positions on either side of the 5′ cut site; then choice of 3′ cut site based on similar choice with base-preference weighting at 3′ end in addition to weighting by empirical length distribution. This generator was run to produce 10,000 footprints from both the actual and a scrambled CxNV1 RdRp sequence, then repeated for a total of 10 runs. The coverage of modeled footprints in each scenario was then compared to the observed coverage using Pearson correlation.

For RNA secondary structure analyses, RNAfold (v2.4.0) from the ViennaRNA package was used to calculate minimum free energy along sliding windows of the stated sizes, with temperature 28 °C (64).

For analyses of footprint context (RNA secondary structure, and amino acid features), a window of adjacent nucleotides/amino acids was first selected. For amino acid analysis, this window included 20 residues upstream of the codon at 2/3 of the distance from the 5′ end of the footprint, roughly corresponding to residues likely to be within the ribosome exit tunnel (28). Features of the sequence within the selected window were then averaged across footprint positions. For analysis probing RNA secondary structure at the edge of footprints, the “local MFE peak” was calculated as (MFE_x_ – average[MFE_(x–win/2)_, MFE_(x+win/2)_]) where MFE_x_ is the MFE at position x and win is the window size, and then compared to the numpy gradient function of the footprint profiles (“footprint boundaries,” emphasizing the plateau edges as steep changes in the coverage).

### Polysome Profiling

CT cell lysates were prepared and separated on 10-50 % m/v sucrose gradients as for ribosome profiling above, except without any RNase digestion step. In addition, the lysate was separated into a control condition or treated with 30 mM EDTA, then incubated on ice for 5 min prior to loading onto sucrose gradients. Fractions were then collected using the Biocomp Gradient Station for the full volume that could be sampled, which excludes the small amount of residual volume of densest sucrose and any pelleted material.

An aliquot of each fraction was then mixed 1:1 with 2× DNA/RNA Shield (Zymo Research) spiked with in vitro transcribed RNA (generated by T7 transcription off a Firefly luciferase (Fluc)-containing plasmid). RNA extraction was performed in 96 well format on a Bravo Automated Liquid Handling Platform (Agilent) using the Quick-DNA/RNA Viral MagBead Kit (Zymo Research) with Proteinase K and DNase steps. RT-qPCR was then performed in 384 well format with Luna Universal Probe One-Step RT-qPCR Kit (NEB), assaying Fluc (primers oHR711/712, FAM probe oHR713), GAPDH (oHR720/721/722), CxNV1 RdRp (oHR729/730/731), and CxNV1 Robin (oHR738/739/740), in triplicate wells for each RNA-target gene combination (Table S9).

Standard analysis was performed on Cq values using empirical amplification efficiencies for each primer/probe set, and the delta-delta Ct method to normalize by Fluc. For each target gene, the relative RNA quantities in each fraction were then normalized across all fractions sampled in the gradient. For each lysate/treatment combination, 2-3 technical replicates were run on separate sucrose gradients and subsequent steps. A total of 2 biological replicates of cell lysates were collected for the non-EDTA condition.

### Data Availability

The complete genome sequence of CxNV1 is available at GenBank under accession numbers MW226855 and MW226856. Raw NGS data have been deposited in the NCBI sequence read archive (SRA) under BioProject PRJNA675022. Raw mass spectrometry data have been deposited at ProteoSAFe (massive.ucsd.edu) under accession number MSV000086532. Supplementary materials available at doi:10.7272/Q6GX48SV.

## ACKNOWLEDGEMENTS

We thank Amy Kistler, Stephen Floor, Jeffrey Hussmann and members of the DeRisi lab for valuable comments; Eric Chow and the UCSF Center for Advanced Technology and the Chan Zuckerberg Biohub for sequencing; Lori Kohlstaedt and the Vincent J. Coates Proteomics/Mass Spectrometry Laboratory at UC Berkeley for mass spectrometry; Kristeene Knopp (formerly DeRisi lab), Valentina Garcia (DeRisi lab), and Wesley Wu (Chan Zuckerberg Biohub) for plasmids; and Aaron Brault (CDC) for the CT and Hsu cell lines.

## Author contributions

H.R. and J.L.D. conceived the study. H.R. designed and performed all experiments and analyzed and interpreted all the data with guidance from J.L.D. K.P. assisted with ribosome profiling and key discussions. M.T.L. and S.S. assisted with sequencing library preparation and RT-qPCR. H.R. wrote the manuscript with input from all authors.

## SUPPLEMENTARY MATERIALS

**Table S1. Sequencing library for parental CT cell line.** Sequenced using paired-end 150nt sequencing on an Illumina NextSeq. Metrics given for IDSeq pipeline processing. QC: quality control filtering. N reads remaining is number of reads after QC filters and host subtraction.

**Table S2. Sequencing libraries for determining 5’ and 3’ ends of CxNV1.** Libraries were sequenced using paired-end 150nt sequencing on an Illumina NextSeq. Adapters were ligated to the molecule type indicated in library name (RNA, or cDNA following reverse transcription), with primers oHR542-545 targeting CxNV1. For all libraries, >97% of reads aligning to CxNV1 aligned in the expected locations at the ends.

**Table S3. Terminal sequences of reads aligning at CxNV1 ends.** All sequences represented at >1% of CxNV1-end-aligning reads shown. Bold indicates sequence with largest fraction. In library RNAlig-542 only 1% of QC-filtered reads aligned to CxNV1 ends (see Table S2).

**Table S4. Variants in CxNV1 detected by mNGS of CT cell line.** The IDSeq pipeline was used to quality filter, subtract host sequences, and de-duplicate PE150 sequencing reads, from which a subset of 999,397 readpairs were then aligned to contigs from *de novo* assembly that corresponded to CxNV1 RdRp and Robin segments. The resulting alignments were analyzed for positions with minimum coverage of 10 reads containing variants at a minimum frequency of 5%. Observed variants are presented here, in coordinates corresponding to complete genomes submitted to Genbank (MW226855 and MW226856). Variants in RdRp were too distant from each other to assess phase. Variants in Robin, despite similar frequencies, were not observed in any consistent combinations (not in phase).

**Table S5. Sequences for CxNV1 mutants.** Locus refers to nucleotide position within full-length viral genome sequence.

**Table S6. CxNV1 peptides by mass spectrometry.** See TableS6.xls. Filtered peptides with hits to CxNV1 proteins from two biological replicates of 1-D LC/MS on CT cell lysate (see methods for details).

**Table S7. Sequencing libraries for ribosome profiling of CT cells.** Libraries were sequenced using single-end 50nt sequencing (SE50) on an Illumina HiSeq, or paired-end 150nt sequencing (PE150) on an Illumina NextSeq. Lib prep indicates library prep method for ribosome footprinting (ribo), or total RNA sequencing (totRNA). Condition indicates pre-treatment before lysis. N raw reads for ribo, or readpairs for totRNA. Duplicate (dup) ratio is calculated as N reads pre-/post-collapse on 100% identity for footprint+UMI (ribo only). CHX: cycloheximide, ribo: ribosome footprinting, totRNA: total RNAseq, Rep: replicate. QC: quality control filtering.

**Table S8. Blast hits for reference host and viral sequences.** See TableS8.xls. Results of searching NCBI nt database using megablast and default parameters. No hits found for CxNV1-Robin.

**Table S9. Primers used in this study.** See TableS9.xls.

**Figure S1. Metagenomic results from IDSeq pipeline processing mNGS data for CT cell line.**

Refers to sequencing library “CT-parental.” Arthropod sequences that were not caught by host filters are assigned non-specifically to a variety of arthropod families, including Aedes, Culex, and Drosophila. No sequences of prokaryotes or non-arthropod eukaryotes were observed. Sequences from 4 viruses were found: an Alphanodavirus (likely all Flock House Virus), Calbertado virus, Phasi Charoen-like phasivirus, and Culex narnavirus 1.

**Figure S2. Hairpin structures at 3’ end of narnavirus RNAs.**

Structures predicted for terminal 30-40 nt of CxNV1 RdRp positive strand encoding RdRp, negative strand with reverse ORF encoding Hypothetical protein, CxNV1 Robin positive strand, and Saccharomyces cerevisiae 20S (Scer20S) narnavirus for comparison (NCBI Accession NC_004051.1). Mean free energy in kcal/mol. Nucleotide color encodes probability of depicted base pairing state. Stop codons of forward ORF shown in red box. Start codons of reverse ORF shown in green box.

**Figure S3. siRNA response to CxNV1-Robin.**

Re-analysis of small RNA sequencing data from Goertz et al. 2019. Top left panel shows abundance of reads in small RNA (Goertz 2019) and in total RNA (this study) mapping to each strand of viral sequences (labeled) and host sequences (unlabeled, <10 rpkm in small RNA). Remaining panels show fragment size distribution of reads mapping to each strand in the uninfected CT cell line (SRR8668667) and in a sample infected with West Nile Virus (WNV) (SRR8668668). Reads in the small RNA data align to CxNV1-Robin (highlighted in yellow) at an abundance comparable to CxNV1-RdRp, with predominantly 21nt fragment length suggestive of an siRNA response.

**Figure S4. Coverage of CxNV1 and CALV from mNGS.**

Coverage shown at each position, using concordantly aligned readpairs from a 1M readpair subset after quality control filtering and host subtraction using the IDseq pipeline. Coverage for negative strand (green) shown at 10-fold magnification relative to positive strand (blue). Position of ORF on forward strand indicated by grey rectangle along x-axis.

**Figure S5. Original gel images for panels shown in Fig 1.**

See FigS5.tif. Lower case letter in brackets above upper left corner of each gel indicates primer pair, according to labeling in main figure (Fig 1). M indicates 1kb+ DNA ladder. Lanes 1 to 8 as in Fig 1C. Red arrow indicates expected band size, centered in each cropped panel on main figure.

**Figure S6. Strand ratio of RNA viruses in wild-caught mosquitoes.**

Analysis of metagenomic NGS libraries prepared from wild-caught mosquitoes in California from Batson et al. 2020 (PRJNA605178). Ratio of reads derived from positive vs. negative strand of each segment of RNA viruses categorized by color. Includes validated viruses and novel viral sequences classified by homology. Each point represents reads from an individual mosquito. CxNV1 was observed in 43 mosquitoes.

**Figure S7. RdRp-dependent persistence of Robin despite residual plasmid detection.**

Amplification of CxNV1-RdRp, CxNV1-Robin, Neo (neomycin on plasmid backbone), or EF1a (host gene) from RNA or DNA isolated from Hsu cells at 3, 6, or 9 weeks after transfection with plasmids indicated in column label. None indicates no plasmid. Empty indicates plasmid backbone with no viral insert. + indicates positive control, either CT cells (for RNA) or plasmid (for DNA). Despite counter-sorting prior to 3 weeks, residual plasmid could be detected as late as 9 weeks post-transfection (amplification of Neo from DNA). CxNV1-Robin is weakly amplified from the DNA fraction in each condition where the plasmid was transfected, but is only detectable in the RNA fraction when co-transfected with RdRp. Representative images shown from n=3 biological replicates. Red arrows indicate rows shown in Fig 2B.

**Figure S8. Original gel images for Fig 2.**

See FigS8.tif. Lanes labeled according to Fig 2 / Fig S7. Lanes marked X not relevant. White text in top left of each image indicates primer target gene. 1kb+ DNA ladder.

**Figure S9. Abundance in footprints vs. total RNA for all reference sequences.**

Data from two replicates of each condition (no drug, CHX) in box plots for footprints (top), total RNA (middle) and density (footprints/totRNA, bottom). Negative strand for host genes not shown because abundance <1rpkm for footprints and total RNA. * for CALV indicates values below axis minimum.

**Figure S10. Expanded gene set for totRNA and footprint coverage.**

Read coverage analysis and visualization as in Figure 3C, except displayed here with negative strand (green) at equal magnification to positive strand (blue). Data shown for CHX-rep1 sample.

**Figure S11. Ratio of positive to negative strand in footprints and totRNA.**

Ratio calculated as number of reads mapping to positive strand divided by reads to negative strand, for persistent viruses detected in CT cells: CALV (positive-sense single-stranded RNA virus), CxNV1, and PCLV (negative-sense single-stranded RNA virus). Data shown for two replicates of each condition (no drug, CHX) in ribosome profiling experiment.

**Figure S12. Expanded gene set for footprint profiles.**

Footprint read mapping and visualization as in Figure 3D. Data shown for two replicates of each condition: no drug (purple), CHX (orange), using normalized values and displayed on a log10 scale. Grey bar indicates cds position on positive strand. Reads mapping to negative strand shown inverted.

**Figure S13. Expanded gene set for footprint length distribution.**

Length distribution analysis and visualization as in Figure 3E (red box indicates identical panels as main figure). Data shown for two replicates of each condition: no drug (purple), CHX (orange), on positive strand and for relevant viruses on negative strand. Parallel analyses for reads before (PRE) and after (POST) collapsing on 100% identity for footprint+UMI. Kolmogorov-Smirnov statistic (*D*) and p-value (p) given for comparison of each gene to EF1a positive strand within the CHX-rep1 sample. With a large D and small p we reject the null hypothesis that both samples are drawn from the same distribution.

**Figure S14. Expanded gene set for periodicity analysis.**

Periodicity analysis and visualization as in Figure 3F. Ribosome densities from 5’ mapping were analyzed using Welch’s method to estimate the power spectral density at each frequency. Frequency of ~0.33 (grey dashed line) corresponds to a period of 3nt. Data shown for two replicates of each condition: no drug (purple), CHX (orange), on positive strand and for relevant viruses on negative strand. EER-contig-7 and EER-contig-40 have the lowest abundance in footprints and totRNA of all host genes (see Figure S9).

**Figure S15. Nucleotide composition in vicinity of footprint edges.**

A) Fraction of each nuclotide at positions ±20nt from 5’ edge (left panel), or ±20nt from 3’ edge (right panel). Position of footprint indicated by grey shading on x-axis. Only footprints within coding region of forward strand used for analysis. Average nucleotide composition for region analyzed given in far right bars. B) log2 of the fold change from average fraction for each nucleotide at designated position (footprint edges) in CxNV1-RdRp, CxNV1-Robin, other viral, or all host genes. Box plot based on host genes.

**Figure S16. Impact of collapsing identical footprint+UMI sequences.**

A) Abundance of footprints mapping to CxNV1-RdRp, CxNV1-Robin, other viral, or host sequences, comparing parallel analyses for reads before (PRE) and after (POST) collapsing on 100% identity for footprint+UMI. Datapoints shown for two replicates of each condition (no drug, CHX). Right panel shows negative strand, where host genes have negligible footprints. B) Data analysis and visualization as in Figure 3D, except that reads POST-collapse are displayed here. Data shown for two replicates of each condition: no drug (purple), CHX (orange), using normalized values and displayed on a log10 scale. Grey bar indicates cds position on positive strand. Reads mapping to negative strand shown inverted. Pearson correlation coefficient (r) between PRE- and POST-collapse coverage profiles for each transcript, averaged across two replicates of each condition.

**Figure S17. Raw count values with 5’ footprint mapping pre/post collapse.**

Footprints mapped at 5’ edge for CxNV1 and two representative host genes in CHX-rep1 sample. Parallel analyses for reads before (PRE, left panels) and after (POST, right panels) collapsing on 100% identity for footprint+UMI. Raw count values shown without normalization. Grey bar indicates cds position on positive strand. Reads mapping to negative strand shown inverted.

**Figure S18. Model of footprint coverage based on nucleotide bias.**

A) Coverage on CxNV1-RdRp of observed footprints in CHX-rep1 sample (left panel), or of 10 runs each with 10k footprints based on nucleotide bias at cut sites and footprint length distribution using the actual CxNV1-RdRp sequence or a scrambled sequence (middle and right panels, respectively). B) Coefficient for Pearson correlation between model-derived profiles and observed footprint profile, for data in (A).

**Figure S19. Analysis of RNA secondary structure.**

A) Mean free energy (MFE) of predicted secondary structures within sliding windows of the indicated size. B) Pearson correlation between MFE (as in (A)) and footprint coverage for CHX-rep1 sample. C) MFE in windows of 15 and 21 nt in vicinity of footprint edges. Footprints aligned at 5′ or 3′ edge (left and right panel for each transcript), covering the region shaded in grey. Mean ± standard error of four samples shown. D) Pearson correlation between local MFE peaks and footprint boundaries across cds length, for CHX-rep1 sample (see Methods for calculation details). E) Examples of footprints vs. potential RNA secondary structure. Top left: magnified view of highest peaks shaded in blue on full profile below. Right: predicted RNA structures for 150-200 nt regions shaded on left, with footprint peaks circled. Arrows indicate 5′ to 3′ direction. Color indicates probability of base pairing for each nucleotide. Note weak hairpins in region 1, vs. stronger hairpins in region 2 which are centered between footprint positions.

**Figure S20. Analysis of nascent chain features.**

Enrichment of amino acids within 20 codons upstream of approximate P-site position, shown as log2 of fold-change relative to abundance in CDS. Boxplot based on host genes, including three outlier datapoints beyond y-axis not indicated by *. Data shown for CHX-rep1 sample. Color of amino acid on x-axis indicates property as follows: non-polar (dark grey), polar uncharged (light green), polar basic (light blue), polar acidic (pink).

**Figure S21. Biological replicate of polysome profiling on CT cells.**

Experiment and analysis as in Figure 3G. Absorbance quantifying total RNA during fractionation of a single gradient shown in lower panel. Relative concentrations of CxNV1-RdRp, CxNV1-Robin, and GAPDH RNA determined by RT-qPCR in upper panel, with n=2 replicates of independent gradients. Grey shading indicates monosome peak.

## REFERENCES

1. Dinan AM, Lukhovitskaya NI, Olendraite I, Firth AE. 2020. A case for a negative-strand coding sequence in a group of positive-sense RNA viruses. Virus Evolution 6.

2. Hillman BI, Cai G. 2013. The family narnaviridae: simplest of RNA viruses. Adv Virus Res 86:149–176.

3. Matsumoto Y, Wickner RB. 1991. Yeast 20 S RNA replicon. Replication intermediates and encoded putative RNA polymerase. J Biol Chem 266:12779–12783.

4. Fujimura T, Solórzano A, Esteban R. 2005. Native replication intermediates of the yeast 20 S RNA virus have a single-stranded RNA backbone. J Biol Chem 280:7398–7406.

5. Charon J, Grigg MJ, Eden J-S, Piera KA, Rana H, William T, Rose K, Davenport MP, Anstey NM, Holmes EC. 2019. Novel RNA viruses associated with Plasmodium vivax in human malaria and Leucocytozoon parasites in avian disease. PLoS Pathog 15:e1008216.

6. Lye L-F, Akopyants NS, Dobson DE, Beverley SM. 2016. A Narnavirus-Like Element from the Trypanosomatid Protozoan Parasite Leptomonas seymouri. Genome Announc 4.

7. Batson J, Dudas G, Haas-Stapleton E, Kistler AL, Li LM, Logan P, Ratnasiri K, Retallack H. 2020. Single mosquito metatranscriptomics recovers mosquito species, blood meal sources, and microbial cargo, including viral dark matter. bioRxiv 2020.02.10.942854.

8. Espino-Vázquez AN, Bermúdez-Barrientos JR, Cabrera-Rangel JF, Córdova-López G, Cardoso-Martínez F, Martínez-Vázquez A, Camarena-Pozos DA, Mondo SJ, Pawlowska TE, Abreu-Goodger C, Partida-Martínez LP. 2020. Narnaviruses: novel players in fungal–bacterial symbioses. 7. The ISME Journal 14:1743–1754.

9. García-Cuéllar MP, Esteban LM, Fujimura T, Rodríguez-Cousiño N, Esteban R. 1995. Yeast Viral 20 S RNA Is Associated with Its Cognate RNA-dependent RNA Polymerase. J Biol Chem 270:20084–20089.

10. Richaud A, Frézal L, Tahan S, Jiang H, Blatter JA, Zhao G, Kaur T, Wang D, Félix M-A. 2019. Vertical transmission in Caenorhabditis nematodes of RNA molecules encoding a viral RNA-dependent RNA polymerase. Proc Natl Acad Sci USA 116:24738–24747.

11. DeRisi JL, Huber G, Kistler A, Retallack H, Wilkinson M, Yllanes D. 2019. An exploration of ambigrammatic sequences in narnaviruses. Scientific Reports 9:1–9.

12. Schlub TE, Holmes EC. 2020. Properties and abundance of overlapping genes in viruses. Virus Evol 6.

13. Brandes N, Linial M. 2016. Gene overlapping and size constraints in the viral world. Biol Direct 11:26.

14. Pavesi A, Vianelli A, Chirico N, Bao Y, Blinkova O, Belshaw R, Firth A, Karlin D. 2018. Overlapping genes and the proteins they encode differ significantly in their sequence composition from non-overlapping genes. PLOS ONE 13:e0202513.

15. Fujimura T, Esteban R. 2007. Interactions of the RNA Polymerase with the Viral Genome at the 5′- and 3′-Ends Contribute to 20S RNA Narnavirus Persistence in Yeast. J Biol Chem 282:19011–19019.

16. Göertz G, Miesen P, Overheul G, van Rij R, van Oers M, Pijlman G. 2019. Mosquito Small RNA Responses to West Nile and Insect-Specific Virus Infections in Aedes and Culex Mosquito Cells. Viruses 11:271.

17. Lin K-C, Chang H-L, Chang R-Y. 2004. Accumulation of a 3′-Terminal Genome Fragment in Japanese Encephalitis Virus-Infected Mammalian and Mosquito Cells. Journal of Virology 78:5133–5138.

18. Ter Horst AM, Nigg JC, Dekker FM, Falk BW. 2019. Endogenous Viral Elements Are Widespread in Arthropod Genomes and Commonly Give Rise to PIWI-Interacting RNAs. J Virol 93.

19. Esteban R, Vega L, Fujimura T. 2005. Launching of the yeast 20 s RNA narnavirus by expressing the genomic or antigenomic viral RNA in vivo. J Biol Chem 280:33725–33734.

20. Hwang J-Y, Buskirk AR. 2017. A ribosome profiling study of mRNA cleavage by the endonuclease RelE. Nucleic Acids Res 45:327–336.

21. Huppertz I, Attig J, D’Ambrogio A, Easton LE, Sibley CR, Sugimoto Y, Tajnik M, König J, Ule J. 2014. iCLIP: Protein–RNA interactions at nucleotide resolution. Methods 65:274–287.

22. Lecanda A, Nilges BS, Sharma P, Nedialkova DD, Schwarz J, Vaquerizas JM, Leidel SA. 2016. Dual randomization of oligonucleotides to reduce the bias in ribosome-profiling libraries. Methods 107:89–97.

23. Dingwall C, Lomonossoff GP, Laskey RA. 1981. High sequence specificity of micrococcal nuclease. Nucleic Acids Res 9:2659–2674.

24. Johnson AG, Grosely R, Petrov AN, Puglisi JD. 2017. Dynamics of IRES-mediated translation. Philos Trans R Soc Lond B Biol Sci 372.

25. Choi J, Grosely R, Prabhakar A, Lapointe CP, Wang J, Puglisi JD. 2018. How Messenger RNA and Nascent Chain Sequences Regulate Translation Elongation. Annu Rev Biochem 87:421–449.

26. Hussmann JA, Patchett S, Johnson A, Sawyer S, Press WH. 2015. Understanding Biases in Ribosome Profiling Experiments Reveals Signatures of Translation Dynamics in Yeast. PLoS Genet 11:e1005732.

27. Lu J, Deutsch C. 2008. Electrostatics in the Ribosomal Tunnel Modulate Chain Elongation Rates. J Mol Biol 384:73–86.

28. Wilson DN, Beckmann R. 2011. The ribosomal tunnel as a functional environment for nascent polypeptide folding and translational stalling. Current Opinion in Structural Biology 21:274–282.

29. Duc KD, Song YS. 2018. The impact of ribosomal interference, codon usage, and exit tunnel interactions on translation elongation rate variation. PLOS Genetics 14:e1007166.

30. Ude S, Lassak J, Starosta AL, Kraxenberger T, Wilson DN, Jung K. 2013. Translation Elongation Factor EF-P Alleviates Ribosome Stalling at Polyproline Stretches. Science 339:82–85.

31. Doerfel LK, Wohlgemuth I, Kothe C, Peske F, Urlaub H, Rodnina MV. 2013. EF-P Is Essential for Rapid Synthesis of Proteins Containing Consecutive Proline Residues. Science 339:85–88.

32. Schlub TE, Buchmann JP, Holmes EC. 2018. A Simple Method to Detect Candidate Overlapping Genes in Viruses Using Single Genome Sequences. Mol Biol Evol 35:2572–2581.

33. Rückert C, Prasad AN, Garcia-Luna SM, Robison A, Grubaugh ND, Weger-Lucarelli J, Ebel GD. 2019. Small RNA responses of Culex mosquitoes and cell lines during acute and persistent virus infection. Insect Biochem Mol Biol 109:13–23.

34. Blair CD, Olson KE. 2015. The Role of RNA Interference (RNAi) in Arbovirus-Vector Interactions. Viruses 7:820–843.

35. Poirier EZ, Goic B, Tomé-Poderti L, Frangeul L, Boussier J, Gausson V, Blanc H, Vallet T, Loyd H, Levi LI, Lanciano S, Baron C, Merkling SH, Lambrechts L, Mirouze M, Carpenter S, Vignuzzi M, Saleh M-C. 2018. Dicer-2-Dependent Generation of Viral DNA from Defective Genomes of RNA Viruses Modulates Antiviral Immunity in Insects. Cell Host Microbe 23:353–365.e8.

36. Wolin SL, Walter P. 1988. Ribosome pausing and stacking during translation of a eukaryotic mRNA. The EMBO Journal 7:3559–3569.

37. Guydosh NR, Green R. 2014. Dom34 Rescues Ribosomes in 3′ Untranslated Regions. Cell 156:950–962.

38. Han P, Shichino Y, Schneider-Poetsch T, Mito M, Hashimoto S, Udagawa T, Kohno K, Yoshida M, Mishima Y, Inada T, Iwasaki S. 2020. Genome-wide Survey of Ribosome Collision. Cell Reports 31:107610.

39. Meydan S, Guydosh NR. 2020. Disome and Trisome Profiling Reveal Genome-wide Targets of Ribosome Quality Control. Mol Cell 79:588–602.e6.

40. Buskirk AR, Green R. 2017. Ribosome pausing, arrest and rescue in bacteria and eukaryotes. Philosophical Transactions of the Royal Society B: Biological Sciences 372:20160183.

41. Ingolia NT, Brar GA, Stern-Ginossar N, Harris MS, Talhouarne GJS, Jackson SE, Wills MR, Weissman JS. 2014. Ribosome profiling reveals pervasive translation outside of annotated protein-coding genes. Cell Rep 8:1365–1379.

42. Irigoyen N, Firth AE, Jones JD, Chung BY-W, Siddell SG, Brierley I. 2016. High-Resolution Analysis of Coronavirus Gene Expression by RNA Sequencing and Ribosome Profiling. PLoS Pathog 12.

43. Machkovech HM, Bloom JD, Subramaniam AR. 2019. Comprehensive profiling of translation initiation in influenza virus infected cells. PLOS Pathogens 15:e1007518.

44. Finkel Y, Mizrahi O, Nachshon A, Weingarten-Gabbay S, Morgenstern D, Yahalom-Ronen Y, Tamir H, Achdout H, Stein D, Israeli O, Beth-Din A, Melamed S, Weiss S, Israely T, Paran N, Schwartz M, Stern-Ginossar N. 2020. The coding capacity of SARS-CoV-2. Nature 1–6.

45. Bercovich-Kinori A, Tai J, Gelbart IA, Shitrit A, Ben-Moshe S, Drori Y, Itzkovitz S, Mandelboim M, Stern-Ginossar N. 2016. A systematic view on influenza induced host shutoff. eLife 5:e18311.

46. Lareau LF, Hite DH, Hogan GJ, Brown PO. 2014. Distinct stages of the translation elongation cycle revealed by sequencing ribosome-protected mRNA fragments. eLife 3:e01257.

47. Main OM, Hardy JL, Reeves WC. 1977. Growth of Arboviruses and Other Viruses in a Continuous Line of Culex Tarsalis Cells. J Med Entomol 14:107–112.

48. Hsu SH, Mao WH, Cross JH. 1970. Establishment of a Line of Cells Derived from Ovarian Tissue of Culex Quinquefasciatus Say. J Med Entomol 7:703–707.

49. Kalantar KL, Carvalho T, de Bourcy CFA, Dimitrov B, Dingle G, Egger R, Han J, Holmes OB, Juan Y-F, King R, Kislyuk A, Lin MF, Mariano M, Morse T, Reynoso LV, Cruz DR, Sheu J, Tang J, Wang J, Zhang MA, Zhong E, Ahyong V, Lay S, Chea S, Bohl JA, Manning JE, Tato CM, DeRisi JL. 2020. IDseq-An open source cloud-based pipeline and analysis service for metagenomic pathogen detection and monitoring. Gigascience 9.

50. Langmead B, Salzberg SL. 2012. Fast gapped-read alignment with Bowtie 2. Nat Methods 9:357–359.

51. Ruby JG, Bellare P, Derisi JL. 2013. PRICE: software for the targeted assembly of components of (Meta) genomic sequence data. G3 (Bethesda) 3:865–880.

52. Zajac P, Islam S, Hochgerner H, Lönnerberg P, Linnarsson S. 2013. Base Preferences in Non-Templated Nucleotide Incorporation by MMLV-Derived Reverse Transcriptases. PLoS One 8.

53. Reuter JS, Mathews DH. 2010. RNAstructure: software for RNA secondary structure prediction and analysis. BMC Bioinformatics 11:129.

54. Tabb DL, McDonald WH, Yates JR. 2002. DTASelect and Contrast: tools for assembling and comparing protein identifications from shotgun proteomics. J Proteome Res 1:21–26.

55. Xu, T, Venable, JD, Kyu Park, S., D. Cociorva, Liao, L., J. Wohlschlegel, J. Hewel,, J.R. Yates III. 2006. ProLuCID, a Fast and Sensitive Tandem Mass Spectra-based Protein Identification Program. Molecular # Cellular Proteomics 5:S174.

56. Cociorva D, L Tabb D, Yates JR. 2007. Validation of tandem mass spectrometry database search results using DTASelect. Curr Protoc Bioinformatics Chapter 13:Unit 13.4.

57. Park SK, Venable JD, Xu T, Yates JR. 2008. A quantitative analysis software tool for mass spectrometry–based proteomics. 4. Nature Methods 5:319–322.

58. McDonald WH, Tabb DL, Sadygov RG, MacCoss MJ, Venable J, Graumann J, Johnson JR, Cociorva D, Yates JR. 2004. MS1, MS2, and SQT-three unified, compact, and easily parsed file formats for the storage of shotgun proteomic spectra and identifications. Rapid Commun Mass Spectrom 18:2162–2168.

59. Peng J, Elias JE, Thoreen CC, Licklider LJ, Gygi SP. 2003. Evaluation of Multidimensional Chromatography Coupled with Tandem Mass Spectrometry (LC/LC−MS/MS) for Large-Scale Protein Analysis: The Yeast Proteome. Journal of Proteome Research 2:43–50.

60. McGlincy NJ, Ingolia NT. 2017. Transcriptome-wide measurement of translation by ribosome profiling. Methods 126:112–129.

61. Dunn JG, Foo CK, Belletier NG, Gavis ER, Weissman JS. 2013. Ribosome profiling reveals pervasive and regulated stop codon readthrough in Drosophila melanogaster. eLife 2:e01179.

62. Gerashchenko MV, Gladyshev VN. 2017. Ribonuclease selection for ribosome profiling. Nucleic Acids Res 45:e6–e6.

63. Dunn JG, Weissman JS. 2016. Plastid: nucleotide-resolution analysis of next-generation sequencing and genomics data. BMC Genomics 17:958.

64. Lorenz R, Bernhart SH, Siederdissen CH zu, Tafer H, Flamm C, Stadler PF, Hofacker IL. 2011. ViennaRNA Package 2.0. 1. Algorithms Mol Biol 6:1–14.

